# Method for assessing coating uniformity of angioplasty balloons coated with poly(lactic-co-glycolic acid) nanoparticles loaded with quercetin

**DOI:** 10.1101/2023.01.18.524614

**Authors:** Allison D. Zieschang, Kevin F. Hoffseth, Tammy R. Dugas, Carlos E. Astete, Dorin Boldor

## Abstract

**Significance:** Drug-coated angioplasty balloons (DCBs) are used to treat peripheral artery disease, and proper dosage depends on coating characteristics like uniformity and number of layers.

**Aim:** Quantify coating uniformity and correlate fluorescence intensity to drug loading for DCBs coated with 5, 10, 15, or 20 layers of poly(lactic-co-glycolic acid) nanoparticles (NPs) entrapped with quercetin.

**Approach:** Images of DCBs were acquired using fluorescence microscopy. Coating uniformity was quantified from histograms and horizontal line profiles, and cracks on the balloons were measured and counted. Fluorescence intensity was correlated with the drug loading of quercetin found from gravimetric analysis coupled with high-performance liquid chromatography (HPLC).

**Results:** Higher numbers of coating layers on DCBs may be associated with less uniform coatings. Cracks in the coating were present on all balloons, and the length of cracks was not significantly different between balloons coated with different numbers of layers or balloons coated with the same number of layers. A strong positive correlation was identified between fluorescence intensity and drug loading.

**Conclusion:** There may be a relationship between the number of NP layers and the uniformity of the coating, but further investigation is needed to confirm this. Fluorescence intensity appears to be a strong predictor of drug loading on DCBs coated with quercetin-entrapped NPs, demonstrating that fluorescent imaging may be a viable alternative to drug release studies.

## 1 Introduction

Peripheral artery disease (PAD) was estimated to affect 236 million people globally as of 2015 and is associated with an increased risk of cardiovascular mortality [1]. PAD is often caused by atherosclerosis, a narrowing of vessels due to plaque buildup. Management of PAD can be medical or surgical. Medical management includes antiplatelet treatment, cholesterol reduction, blood pressure control, and lifestyle changes such as smoking cessation, diet changes, and increased physical activity. In cases where medical management is insufficient, treatment often consists of a percutaneous transluminal balloon angioplasty (PTA) which may be followed by stenting. Balloon angioplasty is performed by inserting a balloon-tipped catheter into an affected vessel. Then, the balloon is inflated to dilate the vessel and achieve patency [2]. Stenting may be performed depending on the length and location of the lesion, as stenting has shown favorable outcomes against restenosis for longer lesions [3]. However, risks of stenting include in-stent restenosis, stent fracture, and thrombosis, making it unfavorable for many PAD lesions [3, 4]. Several common PAD lesion sites, including the superficial femoral artery and the contiguous popliteal artery, are in anatomical areas subject to complex mechanical forces. These forces increase the risks associated with stenting, so angioplasty without stenting is often preferred if possible in PAD lesions [3, 5].

Restenosis, a re-narrowing of vessels that were previously dilated by angioplasty or stenting, is a common complication of endovascular treatment [6, 7]. The inflation of angioplasty balloons causes intimal denuding and stretching vascular injuries; this leads to a cascade of physiological events, including inflammation, cell proliferation, and ECM production, which contribute to intimal hyperplasia [8, 9].

Using nanoparticles (NPs) as a coating for angioplasty balloons shows promising potential for protection against restenosis [6, 10]. NPs on drug-coated angioplasty balloons (DCBs) facilitate the delivery of antiproliferative drugs to prevent the formation of a new lesion, improving patient outcomes of balloon angioplasty [11]. Three foundational components of DCBs are a semi-compliant angioplasty balloon, an antiproliferative drug, and a drug carrier [11]. Paclitaxel-coated angioplasty balloons are approved by the FDA, but a recent meta-analysis identified an increased mortality rate starting two years after treatment for patients treated with paclitaxel-coated balloons and stents compared to control devices [12]. Other concerns related to paclitaxel-coated balloons are systemic toxicity [13], premature release of the drug before reaching the lesion site [13], and delayed re-endothelization [14–16]. The shortcomings of paclitaxel-coated balloons demonstrate the need for other antiproliferative compounds to be investigated for use in DCBs. Quercetin, a flavonoid, has shown promising potential for use in DCBs in vivo and in vitro. It has low toxicity, reduces vascular smooth muscle cell proliferation, and promotes re-endothelization [13, 17, 18]. Quercetin is naturally fluorescent, so DCBs coated with quercetin can be imaged using fluorescence microscopy [19]. Addidionally, drug quantification of quercetin in vivo could potentially be assessed using fluorescence imaging, a sensitive and non-invasive tool [20].

As new drug coatings are developed for DCBs, it is important to investigate the uniformity of the coating, as it is related to drug release [18]. Optimal drug dosage is paramount to ensuring that patients receive the best possible care, and it depends on the characteristics of the drug, patient factors, and disease state [21]. In the case of drugs delivered via NPs, the effective dose of NPs, or the amount of NPs that encounter cells, depends on NP characteristics, core particle and surface chemistry, and the behavior of NPs in vivo [22–24]. Previous research established that the distribution of the drug phase on drug-eluting stents (DES) could impact drug elution kinetics [25]. Although DCBs release drugs over a shorter period of time than DESs, drug distribution is still expected to affect the elution kinetics. Due to the complex nature of NP dosage, quantifying the uniformity of NP-coated surfaces will provide helpful information regarding the quantity of NPs and the surface area free to react with the body’s tissues and fluids. This information will aid in applying current knowledge regarding NP dosing to novel research on DCBs and other NP-coated implants.

## 2 Objectives

This study aims to quantify NP distribution on drug-coated angioplasty balloons and evaluate how the number of NP layers on a DCB affects the uniformity of particle distribution and drug loading. The uniformity of NPs on angioplasty balloons plays a role in determining the optimum number of coating layers. A more uniform distribution on the balloon would translate into a better transfer of drugs onto the blood vessel. Surface irregularities like cracks in the NP coating alter the amount of drug on the surface and could lead to sub-optimal levels of the drug being delivered to the patient. Additionally, this study aims to provide a framework for evaluating the surface uniformity and drug loading of DCBs using non-destructive imaging techniques like fluorescence microscopy. We tested the utility of fluorescence microscopy for comparing coating uniformity between balloons coated with different numbers of layers. This approach could also be applied to compare balloons coated with different drugs or coating systems, and it could serve as a quality control measure in the manufacturing process to monitor the quality of NP coatings. We also correlated fluorescence intensity to drug loading to identify if an equation can accurately predict the amount of drug on a balloon from the fluorescence intensity. This section of of the study was a retrospective analysis of data from our group’s previous work {Craciun, 2022 #105}.

## 3 Materials and Methods

### 3.1 Balloon and NP Coating

Angioplasty balloons were custom ordered from Interplex Medical, LLC (Milford, OH). They featured a 1 mm x 13.9 cm balloon catheter, a 2.7 FR polycarbonate Luer fitting, and a 1.25 mm x 10 mm PET over-the-wire balloon. Eight balloons were coated ultrasonically with poly(lactic-co-glycolic) acid NPs entrapped with quercetin at Sono-Tek (Milton, NY) using a Sono-Tek 48 kHz Accumist nozzle. The coating sample was drawn into a 10 mL syringe and affixed to a MediCoat BCC coating system where it reached room temperature. This system was interfaced with a 3-axis XYZ Gantry System (500mm x 500mm x 100 mm), a rotator, and appropriate hardware to secure the balloon and catheter. The balloons were inflated, secured to the hardware, and coated using 5, 10, 15, or 20 layers with n=2 replicates per coating number. Balloons were named by denoting the number of layers with the letter “L” followed by the number of layers of NPs, and denoting the sample of a particular layer number with the letter “S” followed by the number 1 or 2 (e.g. L5-S1).

### 3.2 Fluorescence microscopy

We measured the uniformity of NPs on the surface of the balloons using fluorescence microscopy. We affixed the balloons to a glass microscope slide using tape and acquired images at 4x magnification with a Cytation 3 Image reader (BioTek Instruments Inc, Winooski, VT). All images were exported using a 16-bit resolution (65536 grey levels) and TIFF format for both the visible range and fluorescent mode. This was conducted according to the procedure previously described by our group {Craciun, 2022 #105}; however, the images were acquired with different parameters.

We collected two sets of images based on different image acquisition parameters: sample-specific and global. The images aquired with sample-specific parmaeters were taken using parameters best suited for each sample to allow for optimal visualization of surface characteristics like cracks and ridges (Table 2). This was done by changing the three acquisition parameters, LED intensity, integration time and gain, to enhance contrast of the image. Coated regions of the balloons appeared bright and cracked regions appeared black, so cracks and other coating abnormalities could be easily identified. We estimated these parameters visually during the image acquisition process according to the perception of an experienced operator. It should be noted that the image for L15-S1 was over-saturated, which may have impacted the data collected from this sample.

**Table 1.** Description and number of cracks for each sample.

| Sample | Number of Cracks | Description |
| --- | --- | --- |
| L5-S1 | 4 | Very fine cracks, primarily in the vertical direction. |
| L5-S2 | 2 | Very fine cracks in the vertical direction. |
| L10-S1 | 4 | One horizontal and vertical crack that appear to be regions where large sections of the coating flaked off the balloon. One additional long horizontal crack and a small diagonal crack. |
| L10-S2 | 4 | One long horizontal crack with a large vertical branch. A few small, isolated vertical cracks. |
| L15-S1 | 14 | Several fine cracks with many branches. Several small, isolated cracks in varying directions. |
| L15-S2 | 11 | Several long horizontal cracks with many branches. A few isolated cracks, mostly in vertical directions. |
| L20-S1 | 6 | Primarily short vertical cracks. One large vertical crack. One large horizontal crack. |
| L20-S2 | 6 | Long horizontal cracks with branches. A few small, isolated cracks in diagonal and vertical directions. |

**Table 2.** Sample-specific image acquisition parameters.

| Sample | LED Intensity | Integration Time (ms) | Camera Gain |
| --- | --- | --- | --- |
| L5-S1 | 3 | 172 | 21 |
| L5-S2 | 3 | 151 | 16 |
| L10-S1 | 2 | 100 | 19 |
| L10-S2 | 3 | 100 | 16 |
| L15-S1 | 3 | 143 | 19 |
| L15-S2 | 3 | 151 | 9 |
| L20-S1 | 3 | 151 | 6 |
| L20-S2 | 5 | 100 | 7 |

We acquired the other set of images using global parameters, so the LED intensity, integration time and gain were the same across all samples (LED intensity=3, integration time=100 ms, camera gain=14). These settings were based on the imaging for a 20-layer balloon (L20-S2). The images acquired with global parmeters for L20-S1 were over-saturated along the edges, but this should not impact the results, given that the analysis was performed along the longitudinal axis of the balloons. The images acquired with global parameters were only used for the retrospective portion of this paper where we correlate fluorescence intensity to drug loading.

The entire balloon was not visible at a 4x magnification, so we collected sequential images that spanned the length of the balloon and then stitched them together. Between seven and nine images were required to capture the length of each balloon. Additionally, L15-S1 exceeded the top and bottom edges, so “top” and “bottom” images were also taken. We used the Stitching plugin from ImageJ to combine the sequential images from each balloon into a complete image {Preibisch, 2009 #40; Schneider, 2012 #122}. For each balloon, we created visible, fluorescent, and superimposed stitched images (**Figure 1**).

**Figure 1.**
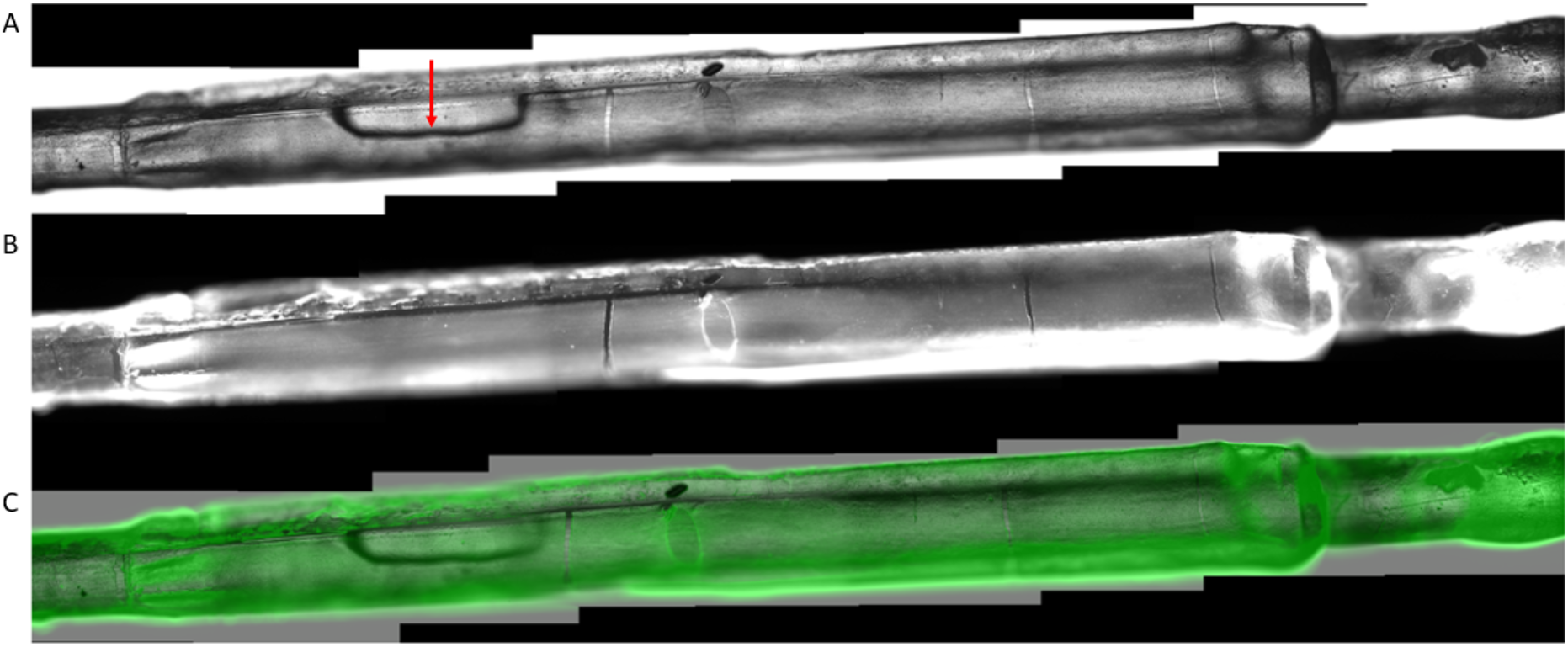
Stitched images acquired with sample-specific parameters for a 10-layer balloon (L10-S2). A-Visible image, B-Fluorescent image, C-Superimposed image with 50% transparency of the fluorescent layer. Note that the longitudinal oval structure in the top and bottom images is the aperture for balloon inflation as indicated by the red arrow in the visible image.

### 3.3 Quantification of NPs

We analyzed the images acquired with sample-specific parameters using ImageJ analysis software by employing two methods to quantify the particle distribution: histograms of rectangular regions of interest (ROIs) and horizontal line profiles. Our images from sample-specific parameters differ from our previous work beacuse they were acquired with unique parameters; therefore, they cannot be used to estimate drug loading based on fluorescence. The image parameters were adjusted for each sample so that none of the images were under or oversaturated which could obscure the visibliity coating features.

The ROIs for the histograms were 100,000 (500×200) pixels in size (1162.79 μm x 465.12 μm), and they captured the middle section of the balloons. We created histograms for a left and right ROI for each unstitched image and used them to evaluate the uniformity of NPs within the ROIs (**Figure 2**). Two histograms were created for each unstitched image, yielding between 14 and 18 histograms per balloon. This was done following the procedure described previously {Craciun, 2022 #105}.

**Figure 2.**
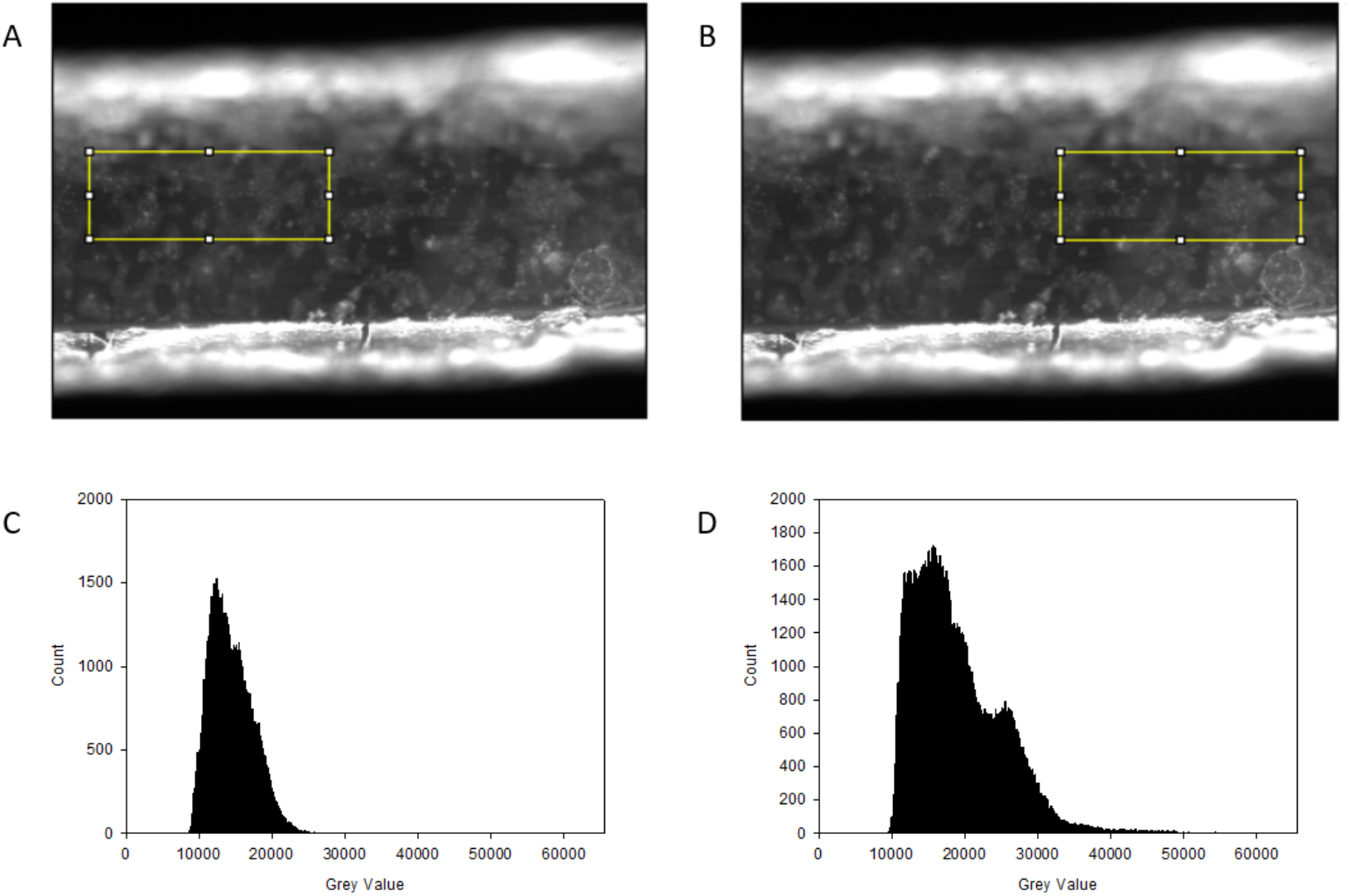
Examples of rectangular selections and corresponding histograms for L5-S2. A-Selection L5-S2-H8, B-Selection L5-S2-H9, C-Histogram L5-S2-H8, D-Histogram L5-S2-H9.

We made horizontal line profiles by creating a 10,000 (800×20) pixel selection parallel to and along the midline of the balloon (1,860.47 μm x 46.51 μm). Then, we generated plot profiles to quantify the grey value at different locations along the selection (**Figure 3** and **Figure 4**). The horizontal line profiles depict the uniformity of NPs along the longitudinal axis; flatter lines indicate more uniform coatings (Figure 5 and Figure 6). Cracks are reflected in the plot profiles as valleys in the grey values. In addition, we investigated cracks in the coatings by counting the number of cracks on each balloon and measuring the length of each crack. To our knowledge, the use of horizontal line profiles is unique to this study.

**Figure 3.**
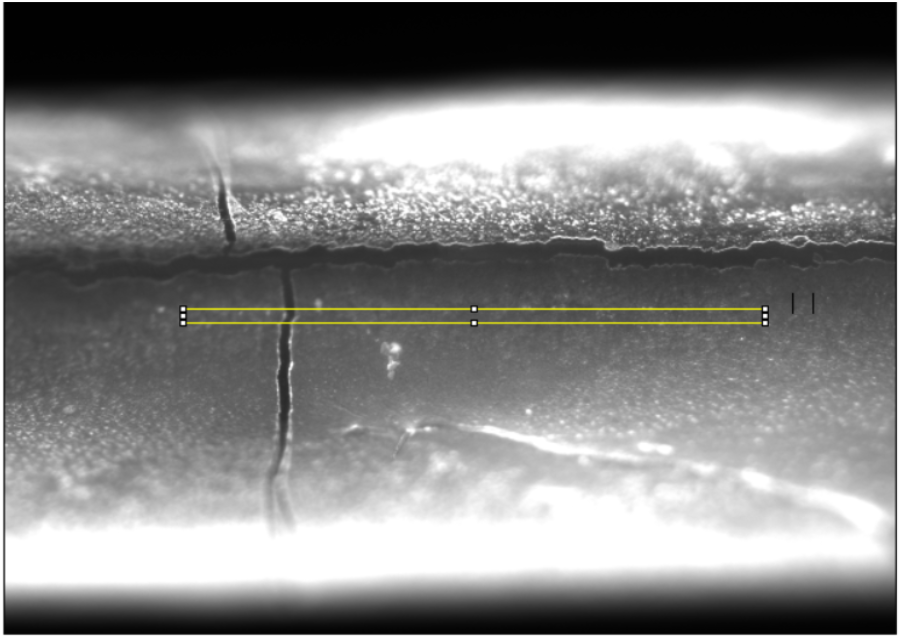
Horizontal line profile selection for L15-S2-HLP6.

**Figure 4.**
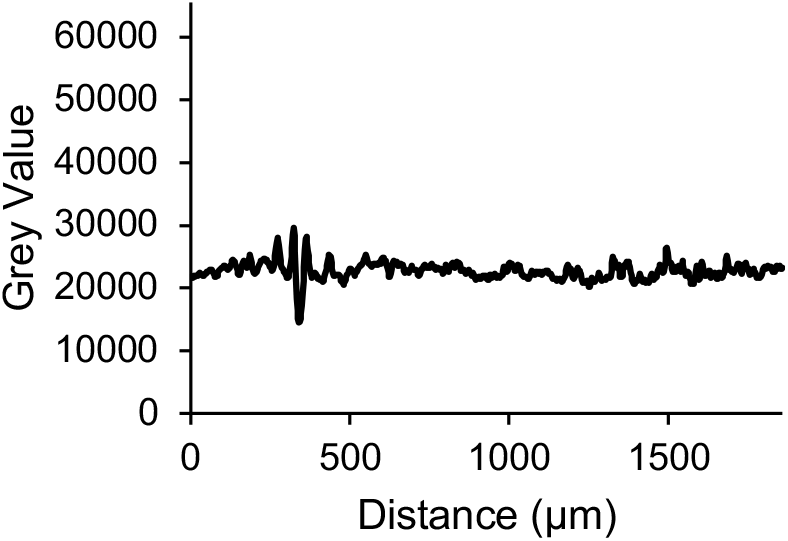
Plot of the grey values for the horizontal line profile from left to right (L15-S2-HLP6).

**Figure 5.**
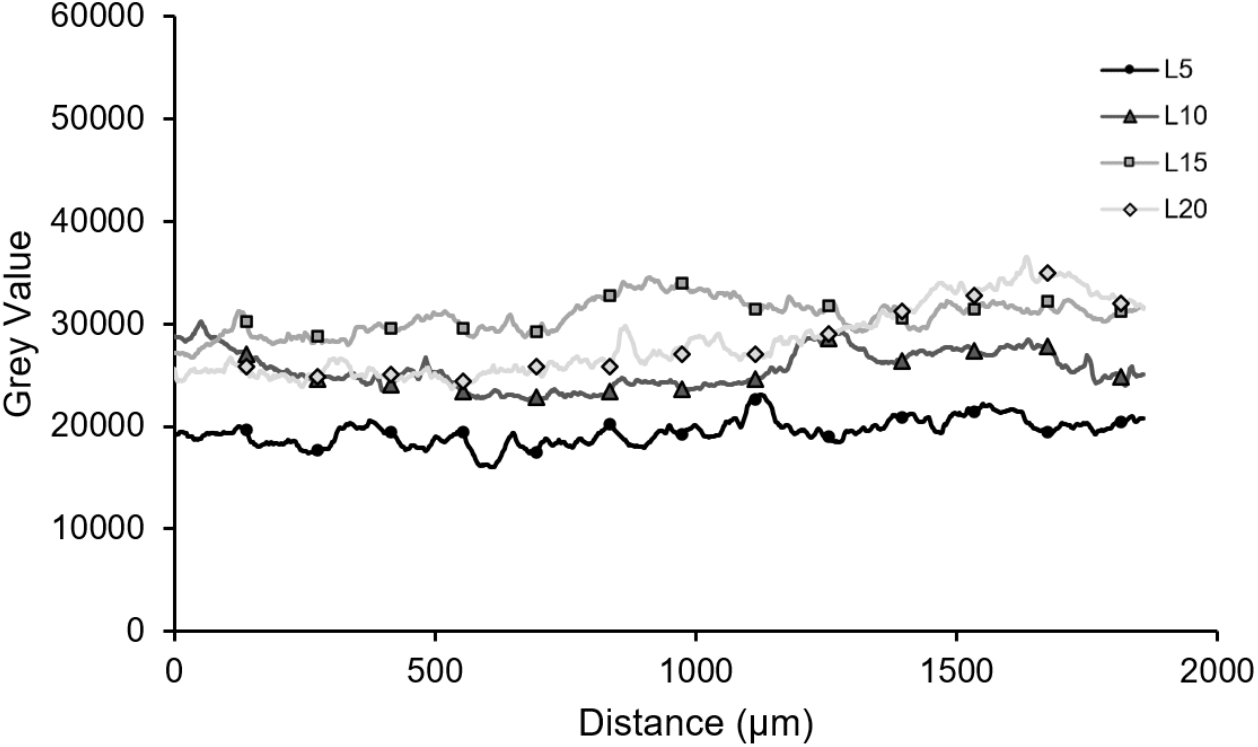
Average grey values of horizontal line profiles by layer number.

**Figure 6.**
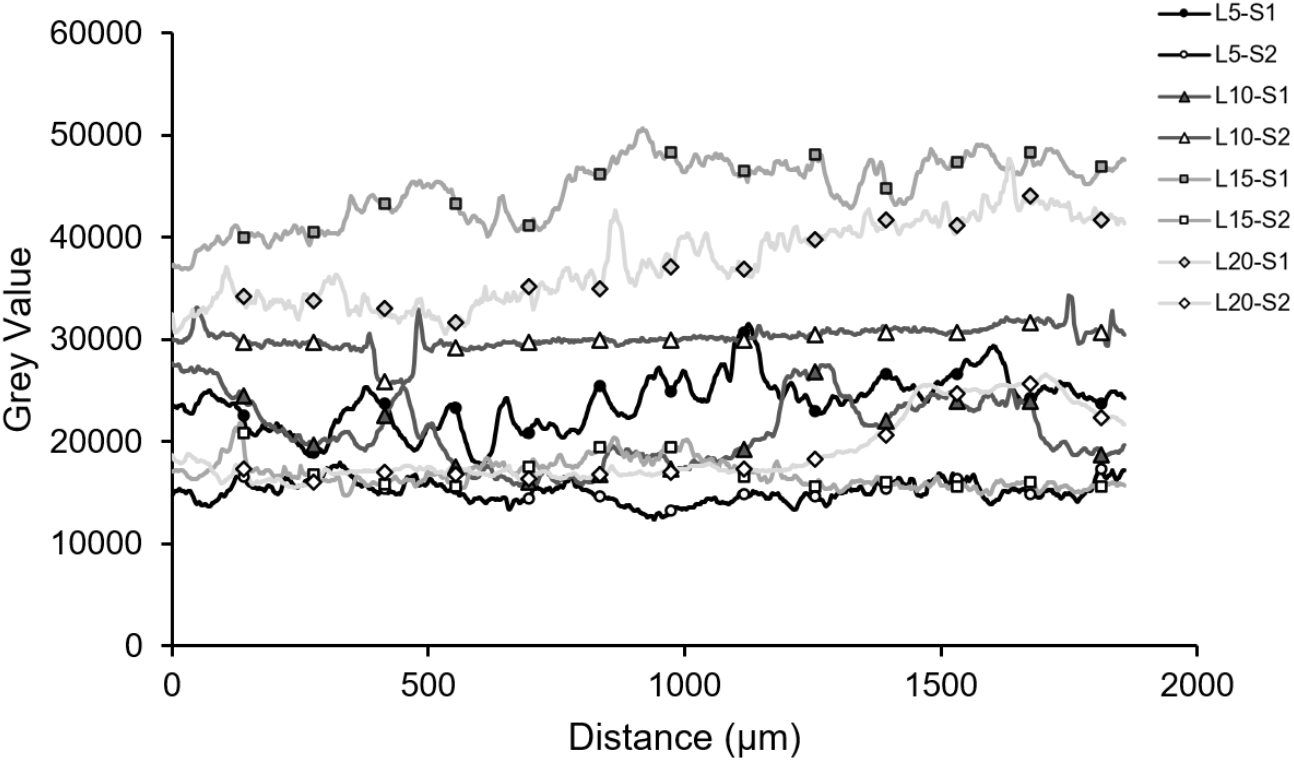
Average grey value of horizontal line profiles by sample.

#### 3.3.1 Uniformity quantification

We quantified coating uniformity using three different criteria. The first measurement of uniformity was based on the standard deviation of each histogram ROI, where higher standard deviations indicate less uniform coating. This approach was previously utilized by our group [18]. The second measurement, also based on our previously published method [18], was found by calculating the percentage of pixels in each histogram ROI within one standard deviation of the mean. Balloons with a larger percentage of pixels within a standard deviation of the mean have a more uniform particle distribution. Lastly, we quantified uniformity along the longitudinal direction of each balloon using the horizontal line profiles. We computed the percent deviation from the mean of the grey values, and smaller percent deviations from the mean indicated a more uniform coating.

#### 3.3.2 Crack analysis

We used the stitched fluorescent images acquired with sample-specific parameters to count and measure cracks in the NP coating. The fluorescent images were used for this analysis because the increased contrast between cracked and coated regions made it easier for the operator to identify and measure cracks. We identified cracks as narrow, dark, line-like segments, with a dramatic change in brightness along the boundaries where the crack meets the coating. Higher numbers of cracks and longer cracks are indicative of sub-optimal coatings.We used the segmented line tool in ImageJ to measure the lengths of cracks. We classified cracks as either branched, vertical, horizontal, or diagonal, and considered cracks with many branches to be a single crack with a combined total length. We then compared the length of cracks between samples and treatments.

#### 3.3.3 Fluorescence intensity and drug loading

Our previous paper discusses the methods for quantifying fluorescence intensity and drug loading [18]. In that work, fluorescence intensity was quantified from the set of images that were acquired with global parameters images. Drug loading was determined by gravimetric analysis and high-performance liquid chromatography (HPLC). In this paper, we preformed a retrospective analysis to correlate the fluorescence intensity and drug loading data that we previously reported {Craciun, 2022 #105}. We did this to determine if fluorescence microscopy is a reliable method for assessing drug loading of DCBs, and we created a linear regression model to predict drug loading from fluorescence intensity.

#### 3.3.4 Statistical testing procedure

We conducted statistical tests for uniformity and cracks using JMP Pro Version 16 (SAS Institute Inc., Cary, NC) to determine whether significant differences exist between different numbers of layers (between treatments) and between samples of the same layer number (within treatments). The histogram analysis was conducted by using the values from each histogram from the unstitched images, so there was a total of 14 to 18 obersvations for each balloon. Likewise, the horizontal line profile analysis was performed by using each of the line profiles constructed on the unstitched images, yielding seven to nine observations per balloon. Cracks were assessed by comparing the number of cracks and the lengths of cracks. We used an alpha value of 0.05, and statistical tests were two-tailed. Before hypothesis testing, we ran a goodness-of-fit test to determine whether the data were normally distributed; since we determined that they were not, we used non-parametric statistical methods for hypothesis testing. We employed a Kruskal-Wallis test to determine whether any significant differences exist between groups. Significant Kruskal-Wallis tests were followed by a non-parametric comparison for each pair using the Wilcoxon method (two-sample Wilcoxon test) to assess the specific differences between balloons coated with different layer numbers and between balloons coated with the same layer numbers.

## 4 Results and Discussion

### 4.1 Uniformity based on images acquired with sample-specific parameters

#### 4.1.1 Histogram: Standard deviation

The standard deviations between balloons with 5 and 15 layers (Z=3.1319, p=0.0017), 5 and 20 layers (Z=4.0309, p<0.0001), 10 and 20 layers (Z=2.3443, p=0.0191), and 15 and 20 layers (Z=2.1480, p=0.0317) were significantly different from each other. Our data do not indicate differences in the standard deviations between the 5- and 10-layer balloons and the 10- and 15-layer balloons. In general, samples with more layers showed greater variance among the ROIs’ standard deviations, probably due to the fact that imperfections in the coating process tend to accumulate as more layers are deposited (**Figure 7Error! Reference source not found.**).

**Figure 7.**
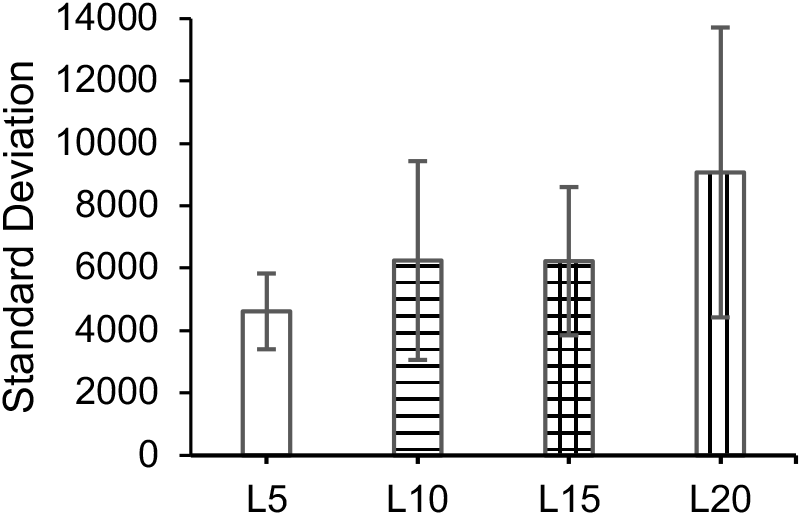
Average standard deviations by loading for histograms, with standard error bars.

*There was a significant difference in the standard deviation between the two samples coated with 5 layers (Z=-3.2270, p=0.0013) and 15 layers (Z=-2.7713, p=0.0056) (**Figure 8**)*. This may result from imperfections in the surface coating or the limited number of replicates per treatment (2 balloons per treatment).

**Figure 8.**
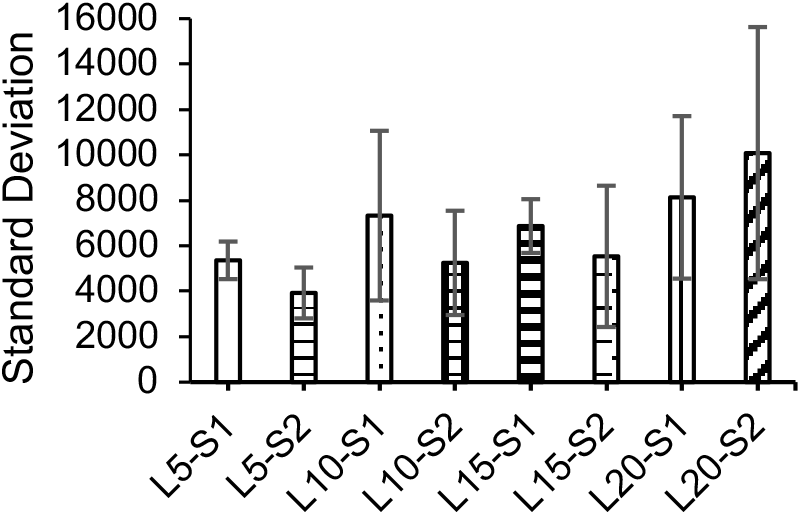
Average standard deviations by sample for histograms, with standard error bars.

#### 4.1.2 Histogram: Percentage of coverage

The percentage of coverage of 5- and 10-layer balloons (Z=2.3981, p=0.0165) and 5- and 20-layer balloons (Z=4.2706, p=0.0111) were statistically different from each other. There is no evidence of differences between any other pair of layer numbers, so the percentage of coverage appears to be independent of layer number. Based on the average percent coverage by layer number, 20-layer balloons have the highest percent coverage, but their coverage was not statistically different from that of balloons coated with 10 or 15 layers (**Figure 9Error! Reference source not found.**). Comparisons of the percentage of coverage of samples with the same layer number did not reveal any significant differences (**Figure 10Error! Reference source not found.**).

**Figure 9.**
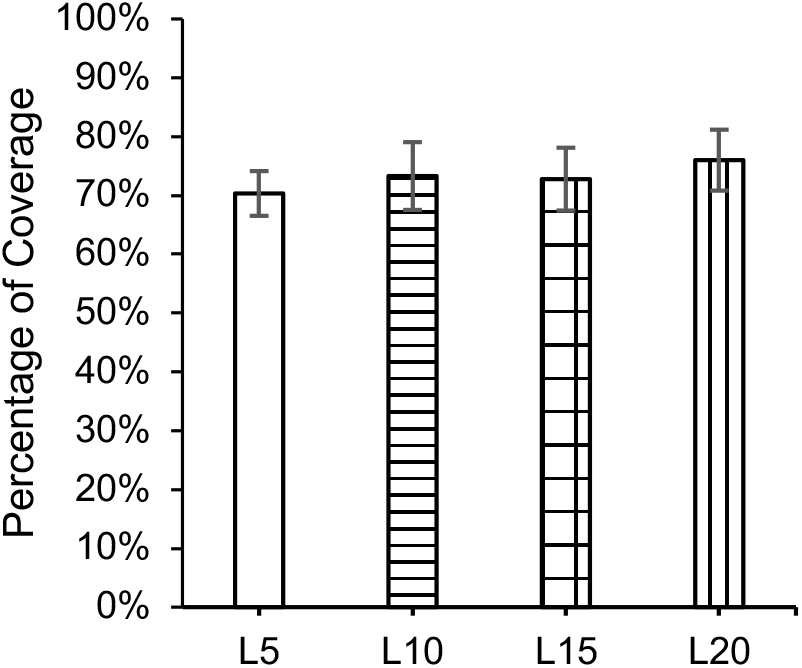
Percent coverage by loading, with standard error bars.

**Figure 10.**
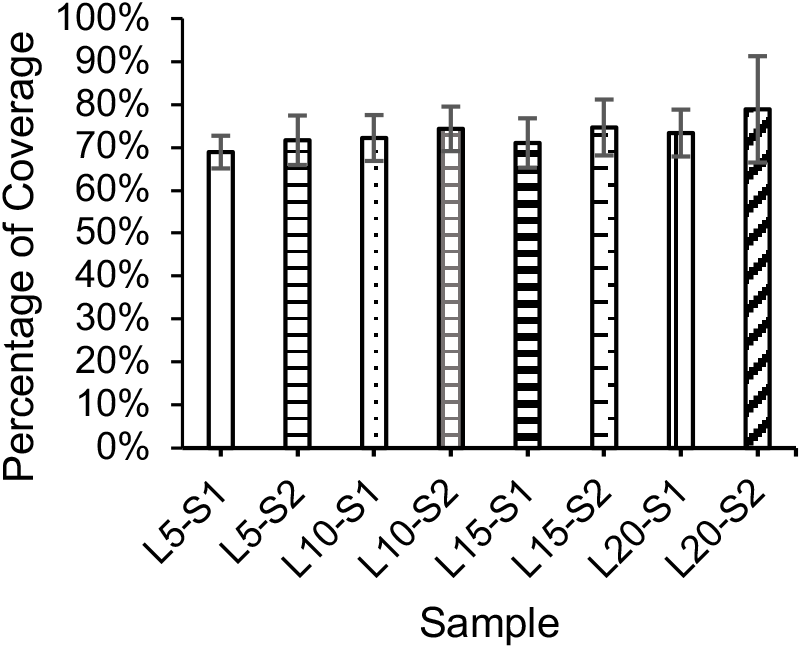
Percentage of coverage by sample, with standard deviation error bars.

To our knowledge, our group is the first to propose using the percentage of pixels within one standard deviation of the mean as a method to assess coating uniformity [18]. Our previous paper proposed a scale to indicate the quality of the coating: <55% poor, 55–60% moderate, 60–70% good, 70–75% very good, 75–80% excellent, and >80% outstanding. However, this approach may be best for comparing differences between balloons rather than determining if a balloon is uniformly coated based on specified cutoff values.

#### 4.1.3 Horizontal line profile: Percent deviation

The percent deviation from the mean gray values was significantly different between 5-layer and 15-layer balloons (Z=-2.6729, p=0.0075). There were no significant differences in the percent deviations between any other pairwise comparison of layer numbers (**Figure 12Error! Reference source not found.**). It is likely that the difference between the 5- and 15-layer balloons are due to the small sample size and factors other than loading, such as cracks induced during the handling of the balloons post-coating.

**Figure 11.**
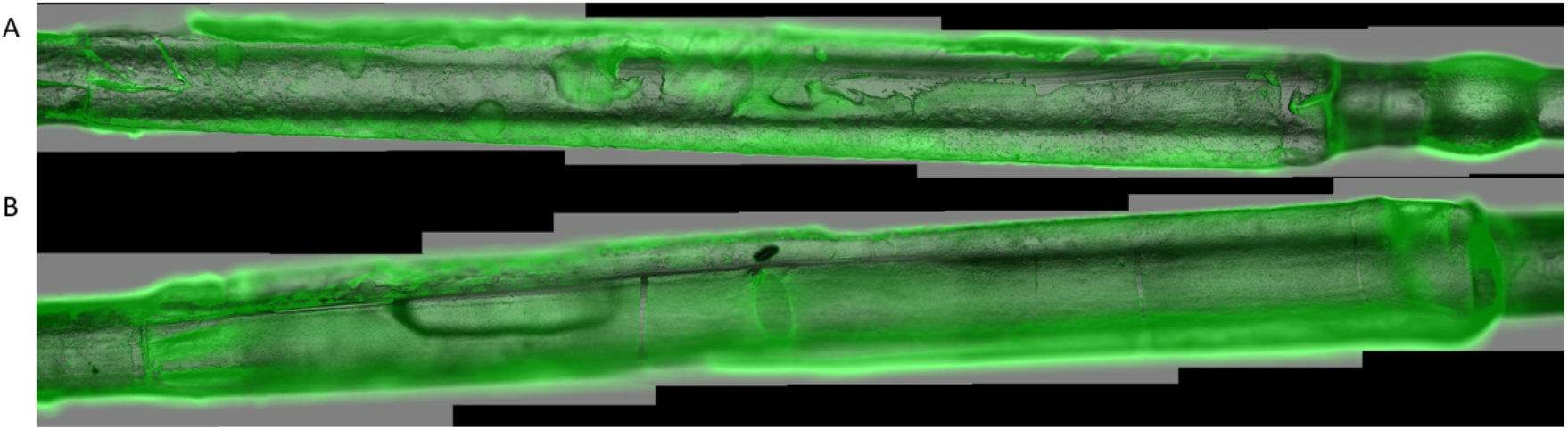
Superimposed images for 10-layer balloons. A-L10-S1, B-L10-S2.

**Figure 12.**
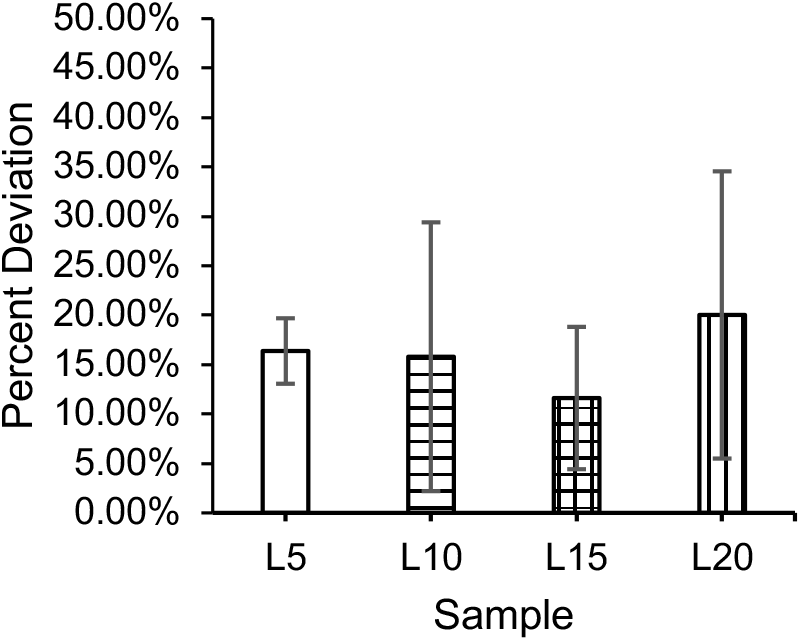
Percent deviation from the mean of gray values by layer number, with standard error bars.

The comparisons between samples of the same layer number indicated a significant difference between the percent deviations of samples coated with 10 layers of NPs (Z=-2.5034, p=0.0123) (**Figure 13**). This result may be due to a poor coating application for L10-S1, which had more visually identifiable anomalies than L10-S2, as shown in **Figure 11**. In general, the average percentage deviation from the mean value was less than 20%, indicating a rather uniform distribution.

**Figure 13.**
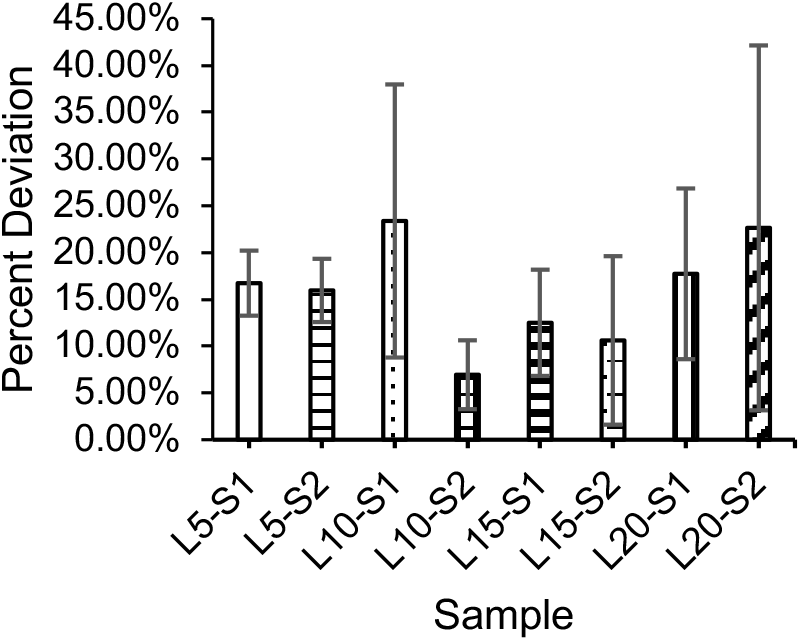
Percent deviation from the mean of gray values by sample, with standard error bars.

#### 4.1.4 Crack Analysis

The balloons coated with 5 layers (L5-S1 and L5-S2) had the fewest cracks, and the cracks were noticeably narrower in the magnified image as qualatively observed. The 15-layer balloons had the highest number of cracks, with many of the cracks having multiple branches. Most cracks occurred in the vertical or horizontal directions, with a few in the diagonal direction. Some cracks contained multiple branches, but the central crack tended to be in the horizontal direction (lengthwise), parallel to the midline of the balloon. The number of cracks with descriptions of the cracking pattern for each sample are detailed in **Table 1**.

The length of cracks varied from 223 μm to 18080 μm across all samples. It is important to note that cracks with multiple branches were analyzed as a single crack with a composite length. The Kruskal-Wallis test comparing the length of cracks between layer numbers found no significant differences in crack length (Figure 14). Additionally, there was no significant differences in the length of cracks between balloons coated with the same number of layers for 10-, 15-, and 20-layer balloons, as found by using the two-sample Wilcoxon test (Figure 15**Error! Reference source not found.**). The differences between the length of cracks for the two samples coated with five layers was not investigated because the number of cracks on L5-S2 was insufficient to preform the analysis.

**Figure 14.**
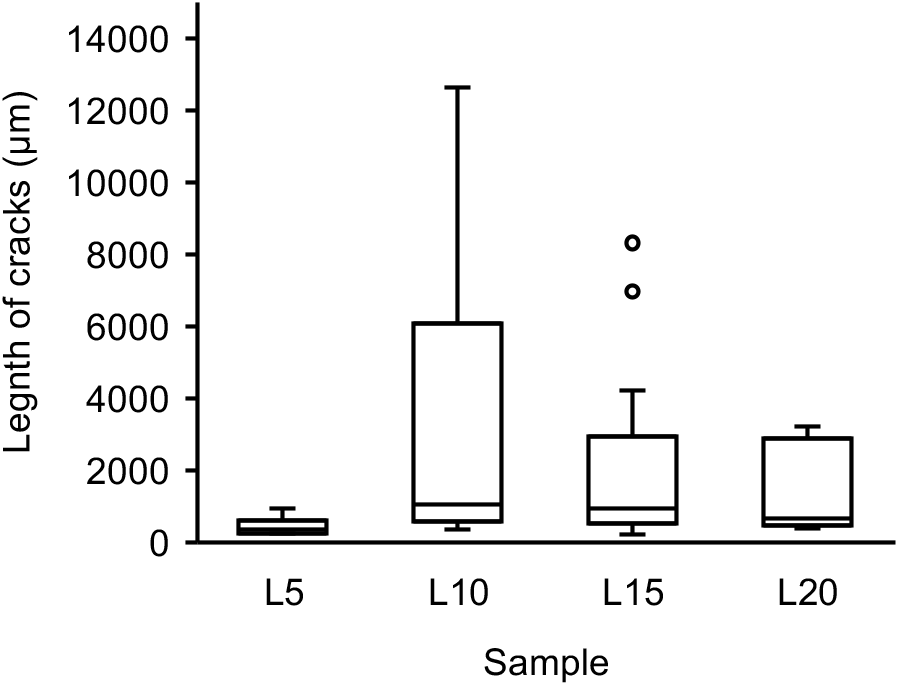
Boxplot of crack lengths by layer number, with standard error bars.

**Figure 15.**
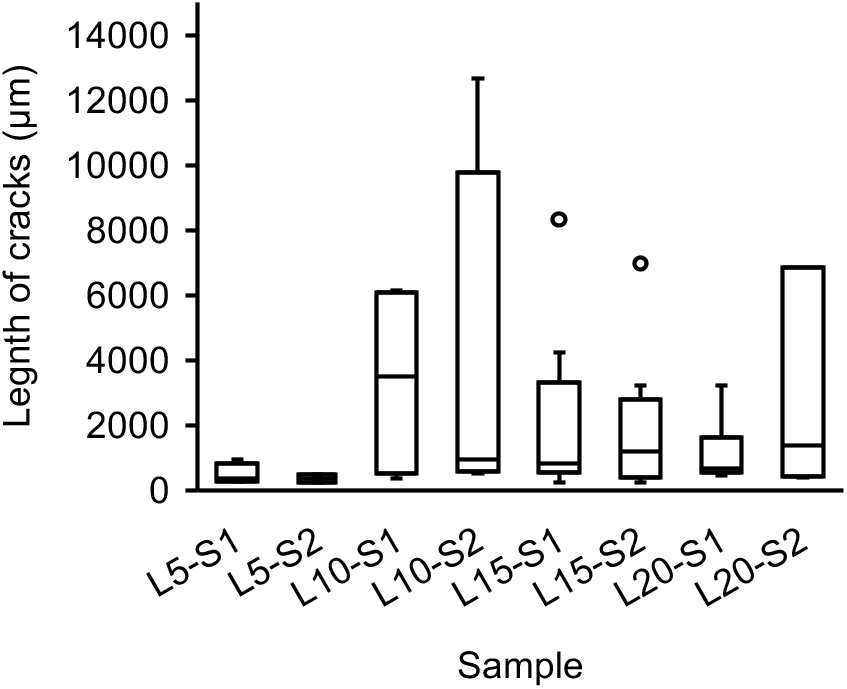
Boxplot of crack lengths by sample, with standard error bars.

The number and length of cracks may be impacted by the coating process in addition to the handling of the balloon post-coating. Affixing the balloons onto the microscope slide could have creased or cracked the coating. Thicker NP coatings are more brittle, and when a brittle coating is applied to a flexible surface like an angioplasty balloon, cracking is likely to occur [18, 28]. Cracks also may have occurred during the drying process and storage of the balloons, and it is possible the imperfections on the base layers of coating may have an accumulating effect as more layers are applied.

### 4.2 Fluorescence intensity and drug loading

We quantified fluorescence intensity as the grey value from histograms created in ImageJ and drug loading as micrograms of quercetin per balloon using HPLC. We found a positive correlation (R = 0.8960) between fluorescence intensity and drug loading. The fitted linear regression model was

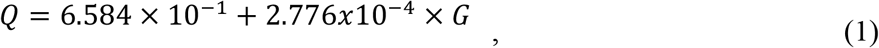

where Q is ug of quercetin on balloon and G is the grey value, and the regression was significant (R^2^ = 0.8028, F[1,6] = 24.43, p = 0.002597) (Figure 16). The significant regression indicates that fluorescence intensity is a good predictor of drug loading, and it demonstrates that calculating the drug loading from fluorescence intensity could be a reliable alternative to preforming drug release studies. This would allow the amount of drug to be calculated without destroying the sample. Future investigation into this correlation would be useful to verify the use of fluorescence intensity to predict drug loading.

**Figure 16.**
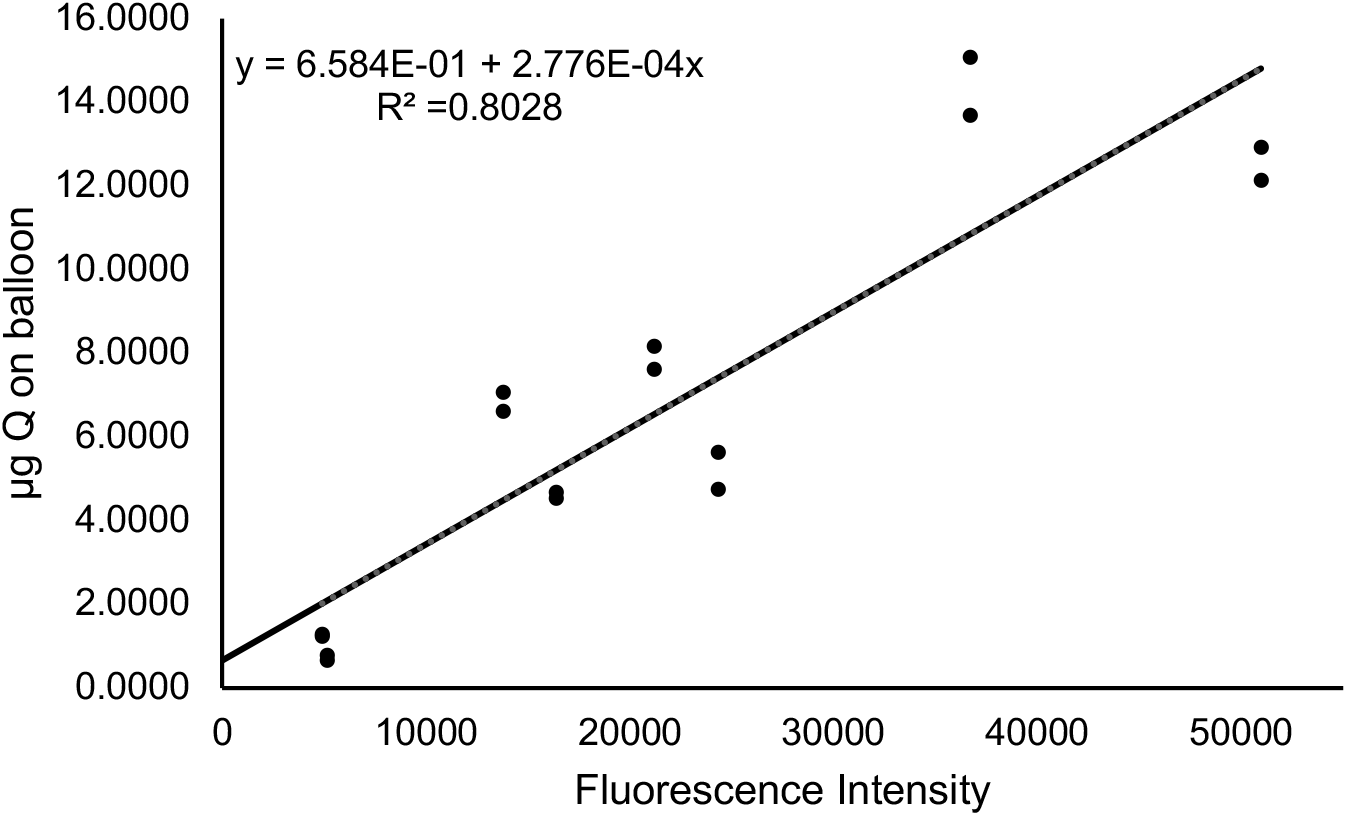
Linear regression plot of fluorescence intensity vs ug of quercetin (Adapted from Craciun, I., et al., *Nanoparticle coatings for controlled release of quercetin from an angioplasty balloon.* PLOS ONE, 2022. 17(8): p. e0268307).

## 5 Conclusion

This approach to quantifying NP uniformity provides insights that can aid in producing uniformly coated DCBs with optimal drug dosage. Additionally, it demonstrates how fluorescence microscopy can be used to investigate and compare the surface uniformity and drug loading of DCBs. Surface uniformity may be influenced by the number of layers of NPs, with higher layer numbers leading to less uniform coatings. This is indicated by the analysis of the standard deviations of the histograms and the percent deviations of the horizontal line profiles, but not the percentage of coverage of the histograms. Future research using larger sample sizes would be necessary to determine how robust this trend is. The percent of coverage within one standard deviation of the mean may be helpful to compare relative differences in coating uniformity, but its use as a statistical approach may be limited because threshold “acceptable” values have not been established.

There were differences in the uniformity between samples coated with the same layer number for 5-, 10-, and 15-layer balloons, but these results were not consistent across all analytical techniques. Crack defects were seen on balloons of all different layer numbers; however, 5-layer balloons had less cracks and shorter lengths as qualatively observed. Defects in the NP coating likely occur during drying, transport, storage, inflation, and handling of the balloons, so special care should be taken when handling DCBs or other NP-coated materials.

We identified a strong positive correlation between fluorescence intensity and drug loading, making fluorescence microscopy a valuable tool for evaluating the surface uniformity and drug loading of DCBs coated with NPs entrapped with quercetin. Due to the non-destructive nature of fluorescence microscopy, it offers advantages over other imaging methods that cannot be coupled with drug release studies. A future investigation comparing the uniformity results from fluorescence microscopy to other established imaging methods, like scanning electron microscopy, would be useful to verify our findings.

## Appendix

## Acknowledgments

This work was funded through a grant award from the National Institutes of Health (R41HL123334) and the Industrial Ties Research Subprogram of the Louisiana Board of Regents Support Fund. Partial support was provided from the USDA NIFA Hatch Program (LAB #94443). This paper is published with the approval of the Director of the Louisiana Agricultural Experiment Station as manuscript # 2023-232-38387.

